# Evolutionary dynamics of mycorrhizal symbiosis in land plant diversification

**DOI:** 10.1101/213090

**Authors:** Frida A.A. Feijen, Rutger A. Vos, Jorinde Nuytinck, Vincent S.F.T. Merckx

## Abstract

Mycorrhizal symbiosis between soil fungi and land plants is one of the most widespread and ecologically important mutualisms on earth. It has long been hypothesized that the Glomeromycotina, the mycorrhizal symbionts of the majority of plants, facilitated colonization of land by plants in the Ordovician. This view was recently challenged by the discovery of mycorrhizal associations with Mucoromycotina in several early diverging lineages of land plants. Utilizing a large, species-level database of plants’ mycorrhizal associations and a Bayesian approach to state transition dynamics we here show that the recruitment of Mucoromycotina is the best supported transition from a non-mycorrhizal state. We further found that transitions between different combinations of either or both of Mucoromycotina and Glomeromycotina occur at high rates and found similar promiscuity among combinations that include either or both of Glomeromycotina and Ascomycota with a nearly fixed association with Basidiomycota. Our results demonstrate that under the most likely scenario symbiosis with Mucoromycotina enabled the establishment of early land plants.

---

Land plants diverged from aquatic algae in the Neoproterozoic as a lineage that would eventually undergo the ecological transition to terrestrial life^1,2^. This transition – a major turning point in the history of life on earth – reshaped the global climate and the biosphere through an increase in atmospheric oxygen levels, carbon fixation, and biotic chemical weathering of rocks^3,4^. Terrestrial life requires plants to extract nutrients and moisture from the substrate. As roots only evolved after the transition to land^5^, initial plant colonization of the terrestrial environment was likely facilitated through interactions with symbiotic fungi where the latter provided inorganic nutrients and water to the host plant and received carbohydrates in return^3,6^.

Mycorrhizal symbiosis is found in over 90% of extant land plant species, and all major lineages of land plants, except for Bryophytes^7,8^. Land plants form associations with members of three different fungal phyla: Mucoromycota, Basidiomycota, and Ascomycota^9,10^. The great majority of land plants associate with arbuscular mycorrhizal fungi from the Mucoromycota subphylum Glomeromycotina, while other types of mycorrhizal associations, such as ectomycorrhiza, ericoid mycorrhiza and orchid mycorrhiza, are formed by fungi of the Basidiomycota or Ascomycota^9^. Fossil evidence suggests that Glomeromycotina have coevolved with land plants for at least 407 Myr, as vesicles, spores, intracellular coils, and arbuscule-like structures resembling extant mycorrhizal infections were found in Rhynie Chert fossils of *Horneophyton lignieri*^11^. Further support for ancient origin of these interactions comes from genomics, as genes involved in the formation of arbuscular mycorrhizal infections are homologs and were acquired in a stepwise manner, with potentiation starting as early as the last common ancestor of Charophytes and Embryophytes^12–14^.

This evidence has led to the wide acceptance of the view that Glomeromycotina were the ancestral mycorrhizal symbionts of land plants^15–16^. The ancestral symbiosis is assumed to have been replaced in several plant lineages by other types of mycorrhizal associations in multiple independent shifts^7^. However, the recent discovery that many members of early diverging lineages of land plants, including liverworts, hornworts, and basal vascular plants, engage in mycorrhizal symbiosis with the Mucoromycota subphylum Mucoromycotina, challenged this hypothesis and suggests that either Mucoromycotina rather than Glomeromycotina could have facilitated terrestrialisation^16^, or that early land plants formed dual Mucoromycotina-Glomeromycotina partnerships^17–20^. After this discovery, Rhynie Chert fossils where re-evaluated, revealing mycorrhizal infections resembling both Glomeromycotina and Mucoromycotina^11^. Moreover, mycorrhiza-formation genes from Mucoromycotina-associated liverworts recover the Glomeromycotina-associated phenotype in a transformed mutant of the angiosperm *Medicago truncatula*, which reveals that the genes required for symbiosis have been conserved among liverworts that associate exclusively with Mucoromycotina as well as higher plants that associate exclusively with Glomeromycotina^13,19^.

Given that Ascomycota, Basidiomycota, Glomeromycotina, and Mucoromycotina diverged prior to the divergence of land plants^21,22^, it is possible to treat different combinations of mycorrhizal association with these phyla as categorical character states on the plant phylogeny and analyse transition dynamics between the states in a Bayesian phylogenetic comparative context. Considering the uncertainty of the evolutionary relationships of early Embryophytes^23,24^, we assessed the probability of all possible combinations of mycorrhizal associations for the most recent common ancestor of land plants.

## Results

We obtained a dataset of 732 species of land plants for which the mycorrhizal fungi have been identified with molecular methods. 45 species were added to represent non-mycorrhizal lineages. We used the plant chloroplast DNA markers *psbA*, *rbcL* and *rps4* to infer phylogenetic relationships between these species. Our estimates of phylogeny correspond well with the prevailing understanding of the systematics of the land plants at least so far as the monophyly of major groups and the relative branching order of these groups under the different rooting scenarios are concerned^25^.

Optimising the observed repertoires of mycorrhizal association as transitioning categorical states on our phylogenetic estimates resulted in a general pattern of phylogenetic conservatism: major plant groups associate quite uniformly with major fungal groups (Figure 1). Our ancestral state reconstructions recover strong support for the presence of mycorrhizal association for the most recent common ancestor of the land plants. However, the particular state for the root was equivocal, showing comparable levels of support for i) an association just with Mucoromycotina, ii) a repertoire comprising both Mucoromycotina and Glomeromycotina; and iii) no mycorrhizal association at all. The relative levels of support, and the inclusion of additionally supported root states, were influenced by different rooting scenarios (Figure S1).

**Figure 1.**
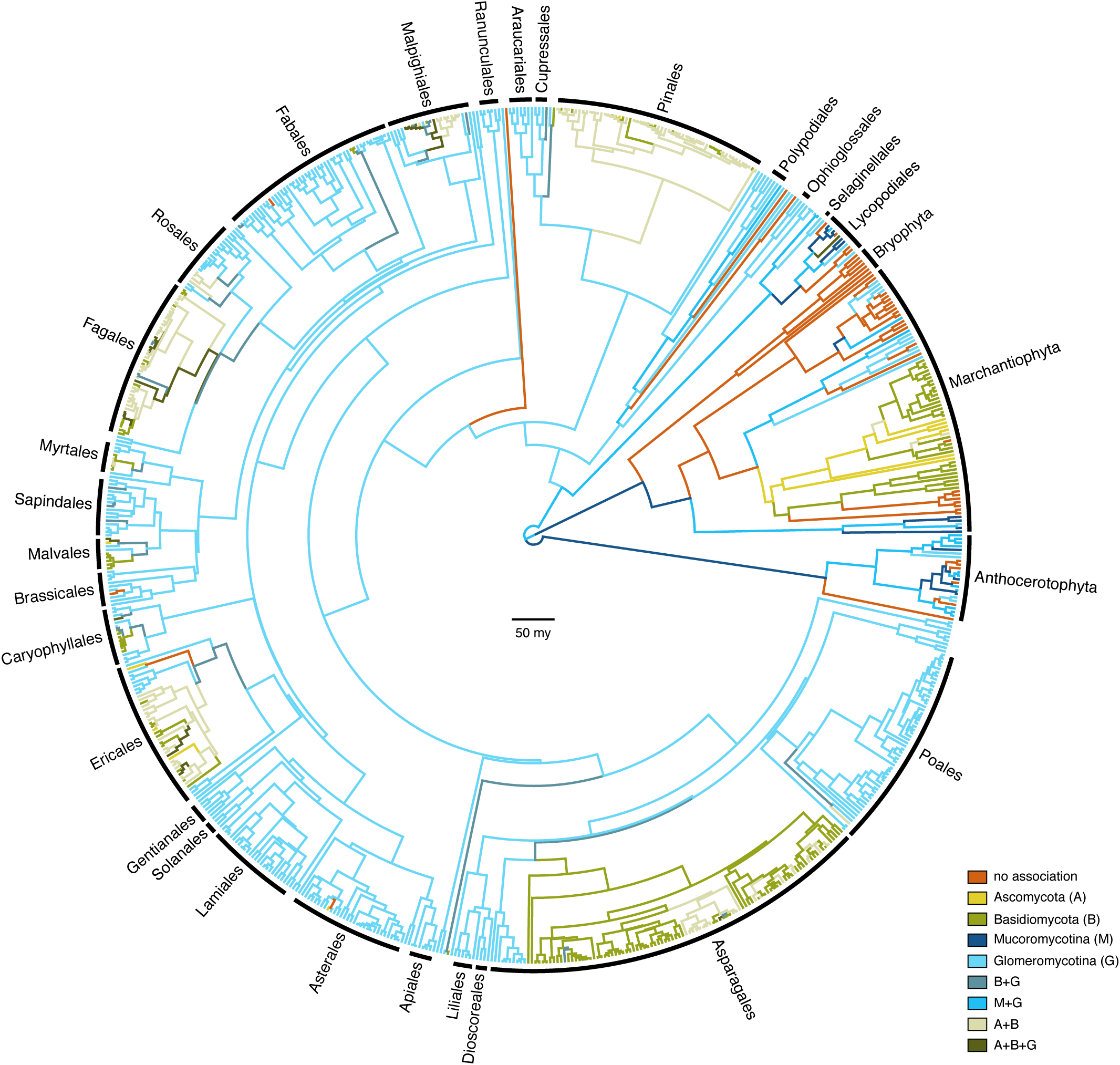
Evolution of mycorrhizal associations in land plants. Chronogram showing the ancestral state reconstructions of mycorrhizal associations in land plants (*n* = 732 species) using a phylogenetic hypothesis in which a clade consisting of liverworts and bryophytes are the sister group of all other land plant species. Branches are coloured according to the most probable state of their ancestral nodes. Main plant lineages are marked with black labels. Branch lengths represent time in million years. Bar is 50 million years.

The pattern of transitions among different repertoires of mycorrhizal association suggests two main paths along which individual associations within a larger repertoire are gained and lost relatively promiscuously (Figure 2). The first of these paths involves Mucoromycotina and Glomeromycotina: the association with Glomeromycotina is added to, and subtracted from, the association with Mucoromycotina at relatively high instantaneous transition rates. The association with Mucoromycotina within a repertoire that spans both is also lost at relatively high rates, but gained at much lower rates, suggesting that the association with Glomeromycotina is relatively more facultative within this repertoire. The second path includes gains and losses of Ascomycota, and losses of Glomeromycotina (but gains less so), at high rates within repertoires in which the association with Basiodiomycota appears more obligate.

**Figure 2.**
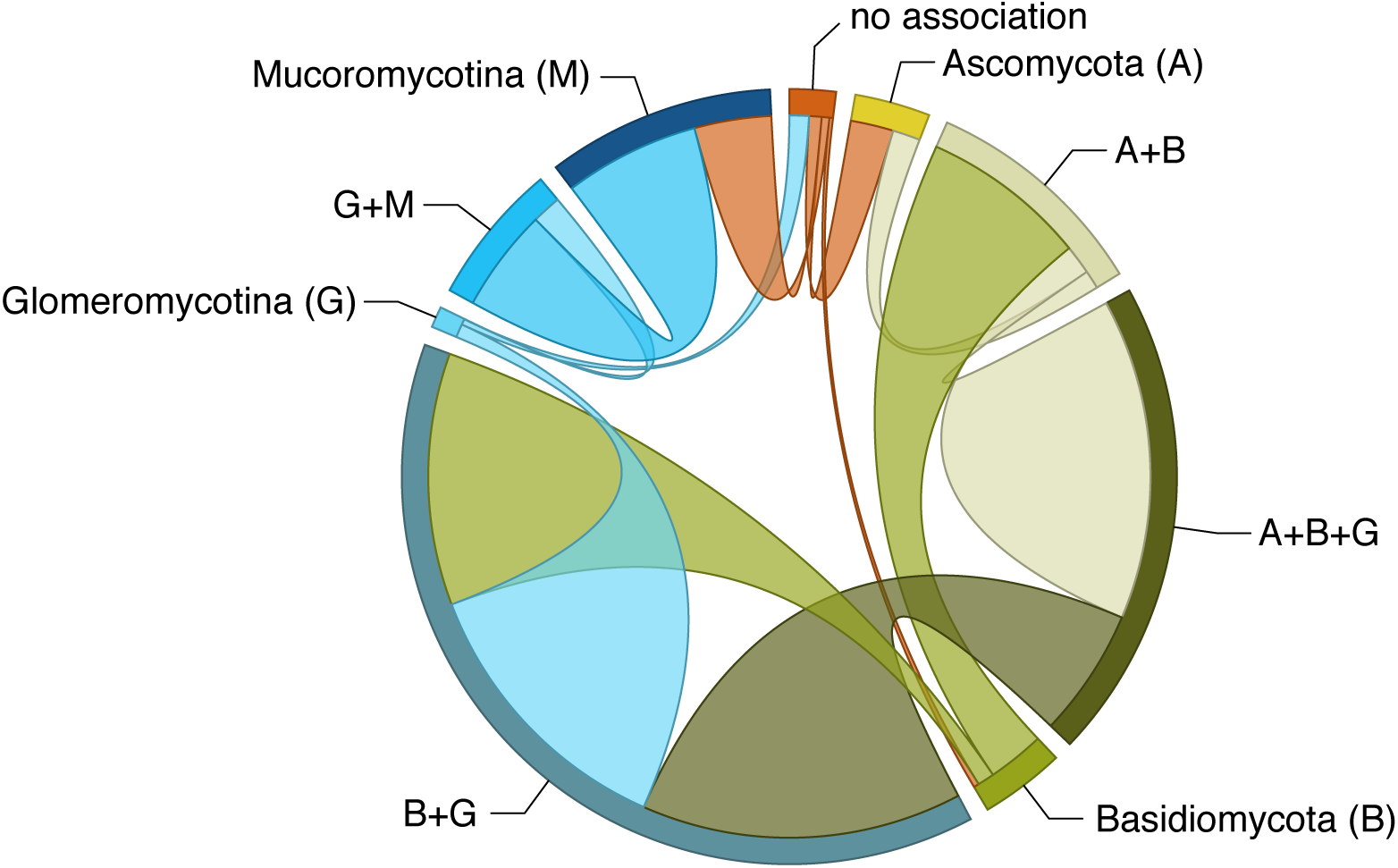
Transitions of mycorrhizal associations in land plant evolution. Frequency of transitions between different repertoires of mycorrhizal association as optimised on our phylogeny (Fig. 1). The band size for each state (labelled next to the bands) represents the number of transitions from that state proportional to the total number of reconstructed transitions; and the width of the ribbons is proportional to the numbers of transitions starting from that state.

Explicit hypothesis testing to quantify which transition away from a state of no mycorrhizal association is best supported prefers Mucoromycotina under all four rooting scenarios: in three out of four, the Bayes Factor (BF) was larger than 10, interpreted as strong support, in the fourth scenario (hornworts sister to all other land plants) the BF was ~8.35, which is generally interpreted as substantial support^26^ (Table S1). Placing the evolution of mycorrhizal associations on a temporal axis in a sliding window analysis (Figure 3) shows Mucoromycotina and Glomeromycotina dominating early associations, while associations that include Basidiomycota and Ascomycota become more pervasive later in land plant evolution.

**Figure 3.**
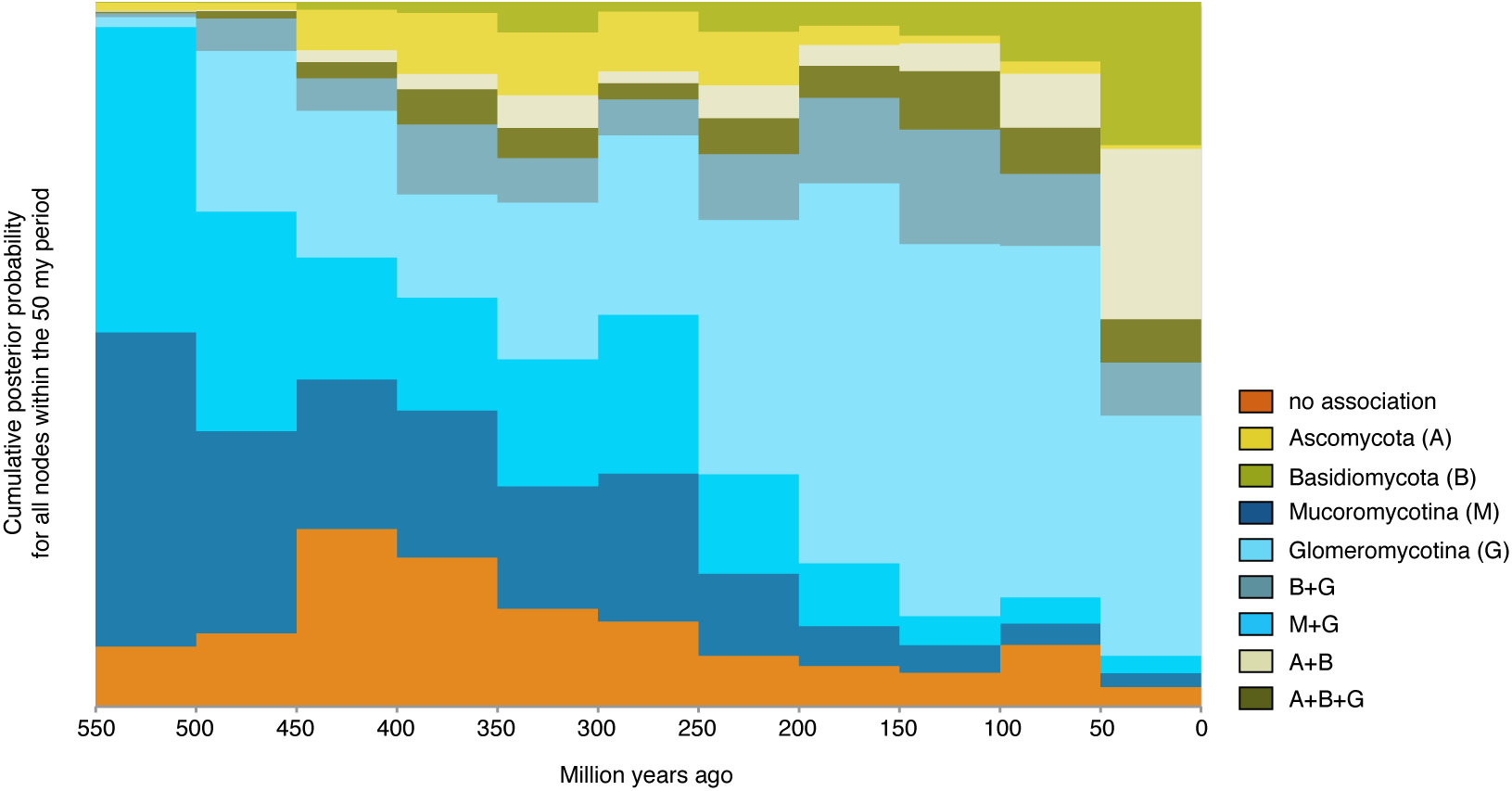
Evolution of mycorrhizal association through time. The proportion of each mycorrhizal state relative to the total number of branches at that particular point in time, sampled at 50 million year intervals on our phylogeny with ancestral state reconstructions (Fig. 1).

## Discussion

For each evaluated scenario of land plant evolution, our results support the hypothesis that the most recent common ancestor of land plants was involved in symbiotic interactions with fungi. This result is in accordance with evidence from the fossil record^11^ and genomics^12–14^. For the small, rootless, leafless plants with simple rhizoid-based absorbing systems that started colonizing the land, the alliance with fungi is hypothesized to have been essential in overcoming major issues of nutrient and water limitation in the absence of existing soils^27,28^. Our analyses suggest that the fungal associates of these earliest land plants most likely belonged to Mucoromycotina, and not Glomeromycotina, as commonly assumed^1,29,30^. An exclusive association with Mucoromycotina for the root of the land plants received the highest support of all possible mycorrhizal repertoires, for all hypotheses of the relationships between the main land plant lineages. Furthermore, our hypothesis tests supported Mucoromycotina over Glomeromycotina as the initial gain for the most recent common ancestor of the land plants. However, our reconstructions also suggest that a repertoire comprising both Mucoromycotina and Glomeromycotina cannot be ruled out, and we find high rates for transitions in which Glomeromycotina are gained and lost in combination with Mucoromycotina (Figure 2), suggesting a versatile scenario for the evolution of association with both groups. Mucoromycotina have been recorded in the rhizoids and roots of extant liverworts, hornworts, lycophytes, and ferns, but except for the liverwort lineage Haplomitriopsida they were mostly found simultaneously with Glomeromycotina^16,17,20^. The association with both fungal lineages was likely also present in the Devonian fossil plant *Horneophyton ligneri*^11^, and Field et al.^18^ speculated that the ability to associate with more than one fungal partner was an ancient strategy that allowed the earliest land plants to occupy highly heterogeneous and dynamic environments. However, this plasticity appears to not be maintained: once association with Mucoromycotina is lost, reversals occur at a low rate (Figure 2) resulting in a predominance of strictly Glomeromycotina associations in extant plants. The scenario presented here is contingent on our current understanding of the early diversification of fungi. Both Mucoromycotina and Glomeromycotina are part of the monophyletic phylum Mucoromycota^10^, and their divergence has been estimated to predate the colonization of land by plants^22^. However, given the large uncertainties of the timing of both events an interaction between early land plants and the common ancestor of Mucoromycota remains a possibility. Under this alternative scenario mycorrhizas formed by Mucoromycotina and Glomeromycotina result from a single evolutionary event within fungi.

From the prevalent association with strictly Glomeromycotina, there have been multiple independent evolutionary shifts towards Ascomycota and Basidiomycota, leading to increasingly prevalent reconstruction of these interactions over the course of plant diversification (Figure 3). Our results suggest that these transitions started with a gain of Basidiomycota, rather than Ascomycota (Figure 2). Subsequent gains of Ascomycota and losses of Glomeromycotina occur at high rates, leading to various association repertoires that include either or both Ascomycota and Basidiomycota. These repertoires are present in several extant land plant lineages. The ability to recruit saprotrophic lineages of wood and litter decaying fungi from among Ascomycota and Basidiomycota into novel mycorrhiza was likely instrumental for plant adaptation to various ecological challenges^5^. For example, for Orchidaceae, the most species-rich lineage of non-arbuscular mycorrhizal plants, the transition from associations with Glomeromycotina to Ascomycota and Basidiomycota is linked to niche expansions and radiations, which in synchrony with the development of specialized pollination syndromes has promoted speciation in the largest family of plants on earth^31,32^. Similarly, the independent evolution of ericoid mycorrhiza in Diapensiaceae and Ericaceae, estimated to date back to the Cretaceous^33,34^, is a potential adaptation to nutrient poor, acidic soils^30^. Also, transitions to ectomycorrhiza independently evolved in various gymnosperm (e.g. Pinaceae, *Gnetum*, *Taxus*) and angiosperm lineages (e.g. Nyctaginaceae, Polygonaceae, Myrtaceae, Malvales, Malpighiales, Fabaceae, Fagales; Figure 1). Parallel to the latter, a shift towards fungi involved in the ectomycorrhizal and ericoid symbiosis has also occurred in liverworts (Figure 1). Although relatively few plant species – mostly trees and shrubs – are ectomycorrhizal, the worldwide importance of the ectomycorrhizal association is considerable, due to its dominance in temperate and boreal forests, and in tropical rainforests in Southeast Asia. Ectomycorrhizal symbioses likely emerged in semi-arid forests dominated by conifers under tropical to subtropical climates and diversified in angiosperms and conifer forests driven by a change to cooler climate during the Cenozoic^35^. Loss of mycorrhizal symbiosis has occurred from all single association states, mostly at relatively low transition rates (Figure 3). These transitions are explained by plant adaptations to either nutrient-rich or extremely nutrient-poor soils, for which the benefits of the symbiosis do not outweigh its costs^36^. However, transition rates towards the non-mycorrhizal state may have been underestimated here, since several non-mycorrhizal angiosperm lineages (all with recent evolutionary origin^37^) have not been included.

Our results portray an evolutionary scenario of evolution of mycorrhizal symbiosis with a prominent role for Mucoromycotina in the early stages of land plant diversification. In most plant lineages, Glomeromycotina, the dominant mycorrhizal symbionts of extant land plants, subsequently replaced Mucoromycotina. Later on, several transitions from Glomeromycotina to various Ascomycota and Basidiomycota lineages have occurred, establishing novel mycorrhizal syndromes, such as orchid, ericoid, and ectomycorrhizas. Our findings demonstrate the importance of Mucoromycotina fungi for our understanding of the early evolution of the mycorrhizal symbiosis. We still know very little about the biology of symbiotic Mucoromycotina. Their presence as mycorrhizal fungi in land plants has been overlooked until recently^16,20^, and it is likely that further screening of land plants will reveal that many more plant taxa are associated with Mucoromycotina.

## Methods

### Data collection

To compile a dataset of plants and their mycorrhizal fungi we searched GenBank for records of Glomeromycotina (at the time of the search ‘Glomeromycota’), Mucoromycotina, Ascomycota and Basidiomycota that had annotations recording the plant host species. Subsequently, for each of these plant host species we conducted a GenBank search and reduced our dataset to all records with an *rbcL* sequence available for the plant host. For the remaining records, we verified mycorrhizal status through literature study and discarded all unconfirmed records from the dataset. We then performed a literature search for plant orders that were not in the dataset as well as for early diverging lineages of land plants.

Because it is difficult to discriminate among mycorrhizas formed by Glomeromycotina and Mucoromycotina by morphological observations, we only included mycorrhizal associations based on DNA identification for these fungi. For lycopods, polypods, hornworts and liverworts, species that were not found to harbour mycorrhizal associations during literature surveys were classified as non-mycorrhizal, although this could be a sampling artefact for some species. Furthermore, mosses and *Nymphaea alba* were included to represent major non-mycorrhizal lineages. The final dataset covers 732 plant species distributed over 78 plant orders. The dataset includes 24 hornworts, 7 mosses, 76 liverworts, 518 angiosperms, 73 gymnosperms, 16 lycopods, and 18 polypods. For these plants species, we found associations with 150 Ascomycota, 305 Basidiomycota, 385 Glomeromycotina, 28 Mucoromycotina and 45 non-mycorrhizal species (Table S2).

DNA sequence data of the plants, including members of the Bryophytes, were obtained from GenBank to reconstruct phylogenetic relationships. For liverworts, hornworts, polypods, and lycopods, we added several species to the dataset to increase taxon sampling, resulting in a total of 759 species for phylogenetic analysis. For 146 species, full or partial chloroplast genomes were available, which we used to extract sequences for *psbA*, *rbcL* and *rps4*. For other species, *rps4* and *psbA* sequences were downloaded where possible, to supplement the *rbcL* dataset. Accession numbers are listed in the supplementary data (Table S3).

### Phylogenetic analysis and divergence dating

For each marker, we aligned the sequences with MAFFT v.7^38^ using the FFT-NS-i Iterative refinement method, and then selected the substitution model with jModelTest 2.1.10^39,40^. For each marker, 3 substitution schemes where tested on a neighbour joining topology, including models with unequal base frequencies, rate heterogeneity and a proportion of invariable sites. The GTR+I+γ model was selected for all partitions using the AIC. We performed divergence dating with BEAST2 v2.3.2^50^ using four fossil calibration points and one age estimate from literature for the crown node of liverworts to date the phylogeny. We selected a uniform distribution for each of the calibration points using the minimum and maximum estimates for these nodes from literature (Table S4). We chose a Yule prior with a uniform birth rate for the analysis, a lognormal relaxed clock model, and estimated the clock rate. We applied the GTR substitution model with a Gamma category count of 4 and estimated shape parameter value of 1.0. The proportion of invariant sites was estimated (initial value 0.01) and the mean substitution rate fixed. We selected an exponential distribution for the prior on the mean substitution rate. To test the effect of different phylogenetic hypotheses^23,42^ for the deep-time relationships of land plants on ancestral state reconstruction, we rooted the consensus tree according to the different hypotheses and applied our divergence dating protocol to each rooted topology (Figure S1). During the MCMC analyses, trace files were updated every 1000 generations, and trees sampled every 10,000 generations, until the effective sample size of major traced parameters exceeded 200 (and all others exceeded 100) using a burn-in of 100 * 10^6^ generations. We thus terminated the runs after, respectively, 374,735,000 generations for ABasal; 354,497,000 generations for ATxMB; 345,720,000 for MBasal; and 374,254,000 for TBasal. We then constructed the maximum clade credibility tree using Tree annotator v2.2.1.

### Comparative analysis and hypothesis tests

In our analysis we assume that the four major fungal groups of which members participate in mycorrhizal associations were already in existence prior to the diversification of land plants^22^. Therefore, we treat each distinct repertoire of associations that land plants form with members of these groups as a discrete state whose evolutionary transition dynamics we modelled subsequent to two additional assumptions. First, because there are qualitative differences between the types of mycorrhizal associations that are formed with some of the different fungal groups (e.g. intracellular versus ectomycorrhizal association), we assumed that the evolutionary adaptations required to enable such associations are not gained (or lost) instantaneously. Hence, we disallowed state shifts that implied multiple, simultaneous gains and losses such that, for example, a change from a state representing a repertoire confined to Glomeromycotina to one confined to Mucoromycotina has to pass through an intermediate state where the repertoire is broadened to include both groups. Second, because the respective adaptations that enable different types of mycorrhizal association are likely subject to evolutionary trade-offs such that repertoires of associations cannot expand infinitely we limited any intermediate states to those we observe in nature. For example, simultaneous association with both Glomeromycotina and Mucoromycotina does occur in our dataset of extant taxa, but complete generalism that includes all fungal groups in a single repertoire does not, which is why we allowed the former, but not the latter, as possible ancestral states.

A convenient side effect of these assumptions was that this limited the number of free parameters in the state transition (*Q*) matrix, which otherwise would have undergone a combinatorial explosion had we included all possible permutations in the repertoires of mycorrhizal association as distinct states, which would have impeded convergence in our analyses. To mitigate such proliferation of potentially unneeded, free parameters further, we performed our analyses using Reversible-Jump MCMC, as implemented in BayesTraits’s ‘multistate’ analysis mode. We ran each of our analyses in triplicate for 10^6^ generations, as initial experimentation had demonstrated reasonable convergence in our data under these settings. In cases where we required estimates of marginal likelihoods, i.e. for hypothesis testing by Bayes factor analysis, we approximated these using a stepping stone sampler that we ran for 100 stones, with 200,000 iterations per stone.

Using this approach, we reconstructed the ancestral states for the four different rootings of our phylogeny. However, although such analyses result in estimates for the posterior distribution of states at any given node (such as the root), they do not necessarily provide the false certainty on which to base a single, unambiguous scenario for the order in which mycorrhizal associations are acquired, especially not when multiple states are reconstructed with similarly large posterior probabilities at deep nodes (as was the case). Given the number of fungal groups and the differences and similarities among these with respect to the types of mycorrhizal associations they participate in, we expected there to be distinct paths along which repertoires of association have evolved. Interrogation and visualisation of the *Q* matrix showed that, broadly, two such paths appear to exist: one where various permutations of association with Glomeromycotina and Mucoromycotina are gained and lost, and another that traverses Ascomycota, Basidiomycota in addition to Glomeromycotina. However, which of these paths was taken first was not yet evident.

We therefore constructed explicit hypothesis tests to distinguish between various plausible scenarios. To do so, in addition to the assumptions affecting the *Q* matrix outlined above, we further constrained our analyses to require the absence of any mycorrhizal association on the root node, and then tested which initial gain was best supported by the data. To quantify this, we estimated the marginal likelihood of the model where the root is constrained to have no association but without any additional constraints on the order in which subsequent associations are acquired (beyond the general assumptions already discussed), and compared this with models where, respectively, each of the initial gains of a single fungal group is disallowed. The logic here is that disallowing the initial shift that best fits the data will result in the marginal likelihood that differs most significantly from the less-constrained model.

Lastly, to place the expansion of repertoires of mycorrhizal association on a temporal axis, we placed the ancestral state reconstructions for the scenario where the root node has no mycorrhizal association in bins of 50 Myr to visualise these in a states-through-time plot (Figure 3). All data and scripts are online available (DOI: https://doi.org/10.5281/zenodo.1037586)

## Acknowledgements

We thank Martin Bidartondo for advice with data compilation, Andrew Meade for help with BayesTraits, and Francis Martin for valuable comments on previous versions of the manuscript.

## Author contributions

V.S.F.T.M initiated and supervised the project. F.A.A.F. compiled the data with input from J.N. F.A.F and R.AV. performed the analyses. All authors wrote the manuscript.

## Competing financial interests

The authors declare no competing financial interests.

## Materials & Correspondence

Correspondence and material requests should be addressed to V.S.F.T.M.

